# Phylogeny and morphology of novel Labyrinthulomycetes in Nova Scotia, Canada

**DOI:** 10.64898/2026.01.15.699578

**Authors:** Jessica Latimer, Eda Ozsan, TJ Goertz, Gurnoor Kaur, Dudley Chung, John M. Archibald

## Abstract

Labyrinthulomycetes are ubiquitous marine decomposers and include the thraustochytrids, a lineage known for their extensive production of polyunsaturated fatty acids and biotechnology applications. Despite historical taxonomic confusion, environmental surveys have begun to expand the known diversity within labyrinthulomycetes. Here we present 18S ribosomal DNA (rDNA) sequences from 17 novel labyrinthulomycetes strains established as axenic cultures from natural samples taken from three locations in Nova Scotia, Canada. We also present light and transmission electron microscopic analysis for three of these isolates. The combination of phylogenetic and morphological analysis suggests that these isolates belong to three different genera, *Thraustochytrium*, *Ulkenia* and *Oblongichytrium*. Our results expand the known genetic diversity of labyrinthulomycetes in Atlantic Canadian waters, contributing new strains to understudied genera for future comparative genomic investigations, including *Ulkenia* and *Oblongichytrium* for which very few genomic resources are available.

## INTRODUCTION

Labyrinthulomycetes are unicellular, heterotrophic protists found ubiquitously in the oceans. Many thrive in nutrient-rich environments abundant in decaying organic matter, such as mangrove forests, sediment layers and river effluents. Most labyrinthulomycetes obtain nutrients by decomposing organic matter; some species have developed parasitic relationships with diatoms (Hassett 2020), flatworms (Schärer et al. 2007) and clams (Geraci-Yee et al. 2021), while others contain photosynthetic endosymbionts (Gomaa et al. 2013), illustrating the diversity of their organismal and ecological interactions. While labyrinthulomycetes were initially classified as fungi due to their complex life cycles and saprotrophy, with the aid of molecular phylogenetics they are now recognized as belonging to the phylum Bigyra, which itself is embedded within the stramenopiles, a large, heterogeneous group of protists and algae (Bennett et al. 2017). Within the class Labyrinthulomycetes are the orders Labyrinthulida, Thraustochytrida, Oblongichytrida and Amphitremida (Bennett et al. 2017).

The class Labyrinthulomycetes has two characteristic ultrastructural features: Golgi-derived scales made from sulfated polysaccharides and an organelle called the bothrosome (historically sometimes called a sagenogenetosome or sagenogen) that produces ectoplasmic networks (Bennett et al. 2017; Iwata et al. 2017). The order Labyrinthulida includes *Labyrinthula* and *Aplanocytrium*, genera that use their ectoplasmic networks for osmotrophic nutrition and as gliding thalli (Bennett et al. 2017). The other major orders, the Thraustochytrida (family Thraustochytriidae: *Aurantiochytrium*, *Botryochytrium*, *Parietichytrium*, *Mucochytrium*, *Schizochytrium*, *Sicyoidochytrium*, *Thraustochytrium* and *Ulkenia*) and Oblongichytrida (family Oblongichytriidae: *Oblongichytrium*) use their bothrosome-derived ectoplasmic nets for anchoring, enzyme secretion and osmotrophic nutrient absorption (Bennett et al. 2017). The Amphitremida (*Archerella*, *Amphitrema*, *Diplophrys*) and one family within Thraustochytrida (the Amphifilidae) do not have true bothrosomes, but instead have pseudostomes and the cytoplasmic extensions take the form of pseudopodia (Pan et al. 2017; Takahashi et al. 2014; Tice et al. 2016).

Due to incongruencies between morphological features and molecular 18S rDNA phylogenies, the class Labyrinthulomycetes has undergone major taxonomic rearrangements (Yokoyama et al. 2007; Yokoyama and Honda 2007). After the erection of new genera in 2007 (Yokoyama et al. 2007; Yokoyama and Honda 2007), namely *Botryochytrium*, *Parietichytrium*, *Sicyoidochytrium*, *Oblongichytrium* and *Aurantiochytrium*, many strains have been reassigned but retain their basionyms. With more 18S rDNA phylogenetic data, the evolutionary relationships between genera have become clearer, but the placement of Oblongichytriidae remains an open question as some studies place it sister to the Labyrinthulida (Takahashi et al. 2014; Gomaa et al. 2013; Anderson and Cavalier-Smith 2012; Yokoyama et al. 2007) and others place it sister to both Labyrinthulida and Thraustochytrida (Pan et al. 2017; Tice et al. 2016; Collado-Mercado et al. 2010; Yokoyama and Honda 2007).

Within labyrinthulomycetes, the thraustochytrids have been intensely studied as they produce large amounts of docosahexaenoic acid (DHA) and are thus of considerable biotechnological interest. Their rapid heterotrophic growth allows for high biomass yield and DHA production, surpassing photo-autotrophic microorganisms limited by light requirements. Prior research has focused on a handful of thraustochytrid genera, including *Aurantiochytrium*, *Schizochytrium* and *Thraustochytrium*, largely due to their capacity for exceptional lipid production. However, genera such as *Ulkenia*, *Oblongichytrium* and *Botryochytrium* are relatively understudied.

With the goal of reducing this imbalance, we are exploring labyrinthulomycete diversity in Nova Scotia, Canada. Here we describe the isolation and characterization of novel strains, including candidate members from neglected lineages, namely the genera *Ulkenia* and *Oblongichytrium*. Our results expand the known genetic and morphological diversity of labyrinthulomycetes and provide a framework for future comparative genomic investigations.

## MATERIALS AND METHODS

### Sample Collection

Sampling was carried out at three sites around Nova Scotia, Canada in May-July of 2024. Samples were collected in sterile Falcon tubes. The pH was assessed with a pH strip, salinity with a refractometer and water temperature with a thermometer.

### Strain Isolation

Isolation was carried out by direct plating and the pollen baiting method (Gaertner 1980). Samples were direct plated on 790 By+ (10 g/L agar, 1 mg/mL yeast extract, 1 mg/mL peptone, 5 mg/mL D-(+)-glucose) plates in artificial seawater (ASW; 24.72 mg/mL NaCl, 0.67 mg/mL KCl, 1.364 mg/mL CaCl_2_ • 2H_2_O, 4.66 mg/mL MgCl_2_ • 6H_2_O, 6.29 mg/mL MgSO_2_ • 7H_2_O, 0.18 mg/mL NaHCO_3_) with penicillin and streptomycin. Plates were incubated at 25 °C and observed daily for growth. The samples were baited with 1 g of autoclaved sterilized pine pollen in 15 mL of ASW for 1 to 2 weeks and then plated again on 790 By+ plates with penicillin and streptomycin. In both cases, colonies were carried forward for single colony isolation before liquid culturing in 790 By+. Liquid cultures were deemed axenic by microscopic inspection and candidate thraustochytrids were identified by morphological appearance with light microscopy.

### DNA Extraction

Cultures were grown in vented Erlenmeyer flasks for 3 days in 790 By+ at 25 °C, on a shaker set to 125 rpm. Cells were pelleted (3220 x g for 10 min) and resuspended in Stan’s extraction buffer (200 mM Tris-HCl pH 7.5, 250 mM NaCl, 25 mM EDTA, 0.5 % SDS). The cell lysate was digested with 0.05 mg/mL RNase A at 37 °C for 15 min and then with 1 mg/mL Proteinase K for 1 h at 55 °C. The samples were extracted twice with equal volumes of 25:24:1 phenol:chloroform:isoamyl alcohol, then twice with equal volumes 24:1 chloroform:isoamyl alcohol. DNA was precipitated overnight at 4°C with 0.1 volumes 3 M sodium acetate, 0.7 volumes isopropanol and 1 μL of GlycoBlue co-precipitant (ThermoFisher cat no. AM9516). The DNA pellet was washed twice with ice cold 95% ethanol and resuspended in Tris-HCl (10 mM, pH 8) buffer.

### Taxonomic Identification

Partial 18S rDNA gene fragments were amplified by polymerase chain reaction (PCR) using primers NSF4/18 (CTGGTTGATYCTGCCAGT) and EukR (TGATCCTTCTGCAGGTTCACCTAC) (https://imr.bio/protocols.html) and 18S-F (GGGAGCCTGAGAGACGGC) and 18S-R (GGCCATGCACCACCACCC) (Mo et al. 2002). Approximately 50 ng of genomic DNA was included in a 25 μL reaction with NEB Q5® High-Fidelity 2X Master Mix and 35 cycles. The annealing temperatures were determined to be 63°C and 72 °C, respectively, with the NEB T_m_ Calculator. PCR products were confirmed on a 1% agarose gel and extracted directly using QIAGEN QIAquick Gel Extraction Kit. The purified PCR products were Sanger sequenced at GENEWIZ by Azenta Life Sciences in the US. New sequences have been deposited in GenBank (accession numbers available upon request).

### Phylogenetic Analysis

Consensus 18S rDNA sequences were built for strains with best BLAST hits to Labyrinthulomycetes using Geneious Prime (2025.2, https://www.geneious.com). Regions with quality scores below q13 were trimmed before sequences were aligned using the Geneious Assembler. A representative set of 52 Labyrinthulomycetes 18S sequences were retrieved from SILVA SSU 138.2 (Chuvochina et al. 2025) as well as outgroup sequences from the stramenopiles *Bacillaria paxillifer* (M87325) and *Ochromonas danica* (M32704). All 18S sequences were aligned using MAFFT-linsi (v7.525) (Katoh and Standley 2013) and trimmed with trimAl (v1.4.rev15) -gappyout functionality (Capella-Gutiérrez et al. 2009). A maximum likelihood phylogenetic tree was inferred using IQTree (v2.3.6) (Minh et al. 2020) with ModelFinderPlus (-m MFP) (Kalyaanamoorthy et al. 2017), 1000 ultrafast bootstrap and 1000 aLRT replicates. The tree was visualized with iTOL (Letunic and Bork 2024).

### Light and Florescence Microscopy

Cultures were seeded on No. 1.5 coverslips (VWR cat no. CA48366-227-1) and grown at 25 °C for 24 to 48 h, then were fixed in PHEM buffer (90 mM PIPES, 37.5 mM HEPES, 15 mM and 3 mM MgCl_2_ • 6 H_2_O) and 4% formaldehyde for 30 min at room temp. Coverslips were washed 4 times with PHEM buffer solution (5 min each), stained with 5 µM Hoechst 33342 prepared in PHEM buffer solution for 10 min at room temp, then washed once in PHEM buffer solution for 5 min. Samples were mounted on microscope slides (Fisher Scientific cat no. 12-550-15) with Prolong Diamond antifade (ThermoFisher cat no. P36965). DIC and fluorescence images were captured using the Axiocam 506c and Axiocam 506m digital cameras, mounted on a Zeiss Axio Imager Z1 widefield microscope equipped with an HXP 120 light source.

### Transmission Electron Microscopy

Cells were pelleted (3220 x g for 10 min) and fixed in 2.5% glutaraldehyde prepared in 1.5X PHEM buffer with 9% sucrose at 4°C for 90 h. Samples were rinsed three times with 0.1M sodium cacodylate buffer, fixed for 2 h with 1% osmium tetroxide, then rinsed with distilled water. Samples were stained overnight at 4°C with 0.25% uranyl acetate, dehydrated using a graduated series of acetone (50%, 70% twice, 95% twice, 100% twice), embedded with Epon Araldite resin (3:1 dried 100% acetone to resin for 3 hours, 1:3 dried 100% acetone to resin overnight, 2x 3 hrs 100% resin) and cured at 60°C for 48 h. Thin (80-100 nm) sections, obtained using a Reichert-Jung Ultracut E ultramicrotome, were placed on 300 mesh copper grids and further stained with 2% aqueous uranyl acetate and lead citrate. Samples were imaged with a Hamamatsu ORCA-HR digital camera attached to a JEOL JEM 1230 microscope at 80kV, or with a Gatan 832 ORIUS® SC1000 digital camera attached to a FEI Tecnai-12 microscope at 80kV.

## RESULTS & DISCUSSION

### Environmental Sampling of Labyrinthulomycetes

We sampled 16 sites from three regions in Nova Scotia, Canada: Boggy Hill (BH), Cape Breton (CB), as well as Cole Harbour near Cow Bay (Cows) and Ira Settle Point (Ira) (**Figure 1**, **Table 1**). A combination of pine pollen baiting and direct plating with antibiotics was used to isolate members of the protist class Labyrinthulomycetes. Pine pollen is a selective bait for thraustochytrids because their ectoplasmic nets penetrate the sporopollenin layer of pine pollen to feed the thallus (Raghukumar 2002; Gupta et al. 2013). Thraustochytrids are known for their ability to decompose plant detritus and many have been isolated from decaying plant matter like mangrove leaves. In a culture starved of other nutrients, thraustochytrids will colonize the pine pollen grains and become enriched.

**Figure 1.**
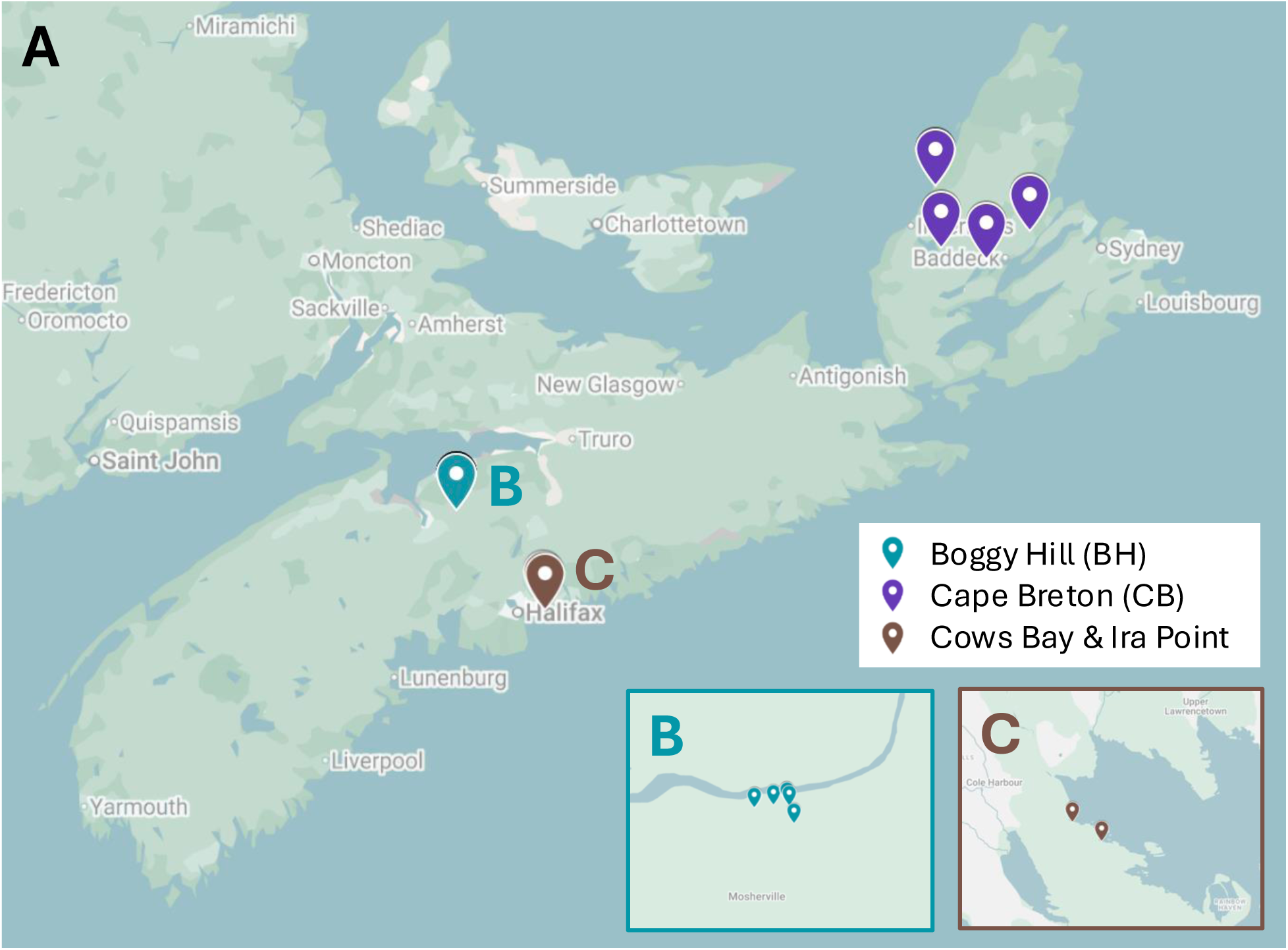
Map of Nova Scotia, Canada with sampling sites indicated. **A.** Boggy Hill (BH1-9) in teal, Cape Breton (CB1-5) in purple, Cole Harbour near Cow Bay (Cows) and Ira Settle Point (Ira) in brown. **B.** Close up of Boggy Hill sites. **C.** Close up of Cow Bay and Ira Settle Point sites in Dartmouth, Nova Scotia.

**Table 1.** Description of 16 Nova Scotian sampling sites and environmental conditions. The main sampling regions are listed Boggy Hill (BH1-9), Cape Breton (CB1-5), Cole Harbour near Cows Bay (Cows) and Ira Settle Point (Ira). The sampling date and coordinates are listed, alongside the water pH, temperature and salinity. The sites in grey did not yield Labyrinthulomycetes.

| Site | Date | Coordinates |  | pH | T (°C) | Salinity |
| --- | --- | --- | --- | --- | --- | --- |
| BH1 | 13-May-24 | 45°4'15" N | 63°57'58" W | 6.5 | 11 | 1.30% |
| BH2 | 13-May-24 | 45°4'15" N | 63°57'58" W | 6.5 | 10 | 1.30% |
| BH3 | 13-May-24 | 45°5'34" N | 64°5'20" W | 6.5 | 10 | 1.30% |
| BH4 | 13-May-24 | 45°4'15" N | 63°58'0" W | 6.5 | 14 | 1.20% |
| BH5 | 13-May-24 | 45°4'14" N | 63°58'0" W | 6.0-6.5 | 14 | 1.40% |
| BH6 | 13-May-24 | 45°4'15" N | 63°57'53" W | 6.0 | 12 | 0.20% |
| BH7 | 13-May-24 | 45°4'15" N | 63°57'54" W | 5.5 | 16 | 0.30% |
| BH8 | 13-May-24 | 45°07' N | 63°97' W | 6.0 | - | 0.20% |
| BH9 | 13-May-24 | 45°4'1" N | 63°57'50" W | 5.5-6.0 | 16 | 0.15% |
| CB1 | 29-Jun-24 | 46°08'35" N | 61°08'39" W | 5 | 18 | 0.10% |
| CB2 | 29-Jun-24 | 46°23'27" N | 61°10'06" W | 7.5 | 15 | 0.20% |
| CB3 | 29-Jun-24 | 46°23'28" N | 61°10'11" W | 8.5 | 16 | 3.80% |
| CB4 | 30-Jun-24 | 46°12'26" N | 60°37'17" W | 7 | 16 | 2% |
| CB5 | 30-Jun-24 | 46°05'39" N | 60°52'25" W | 6 | 17 | 1.20% |
| Cows | 02-Jul-24 | 44°39'43" N | 63°27'04" W | 7 | 18 | 4% |
| Ira | 02-Jul-24 | 44°39'56" N | 63°27'31" W | 5.3 | 18 | 2.20% |

Twenty-seven clonal cultures were established, 21 from baiting with pollen grains of wild lodgepole pine (*Pinus contorta*) and six from direct plating onto solid medium containing antibiotics. Clonality was inferred by light microscopy where the cells appeared globose or sub-globose. Taxonomic identity was confirmed using PCR amplification and direct sequencing of 18S ribosomal DNA (rDNA) followed by BLASTn searches of partial 18S rDNA amplicons against the NCBI nucleotide database. Of the 27 cultures, ten were identified as off target organisms, primarily fungi (data not shown). The pine pollen baiting and direct plating each yielded five off target cultures, but pollen baiting was more fruitful with 16 of the 17 cultures established using this method identified as Labyrinthulomycetes (**Table 2**). Direct plating only yielded one putative labyrinthulomycete, designated clone CB4P3. The best BLAST hits for 18S rDNA suggested our 17 labyrinthulomycetes isolates are primarily from the genera *Thraustochytrium*, *Ulkenia* and *Oblongichytrium*.

**Table 2.**
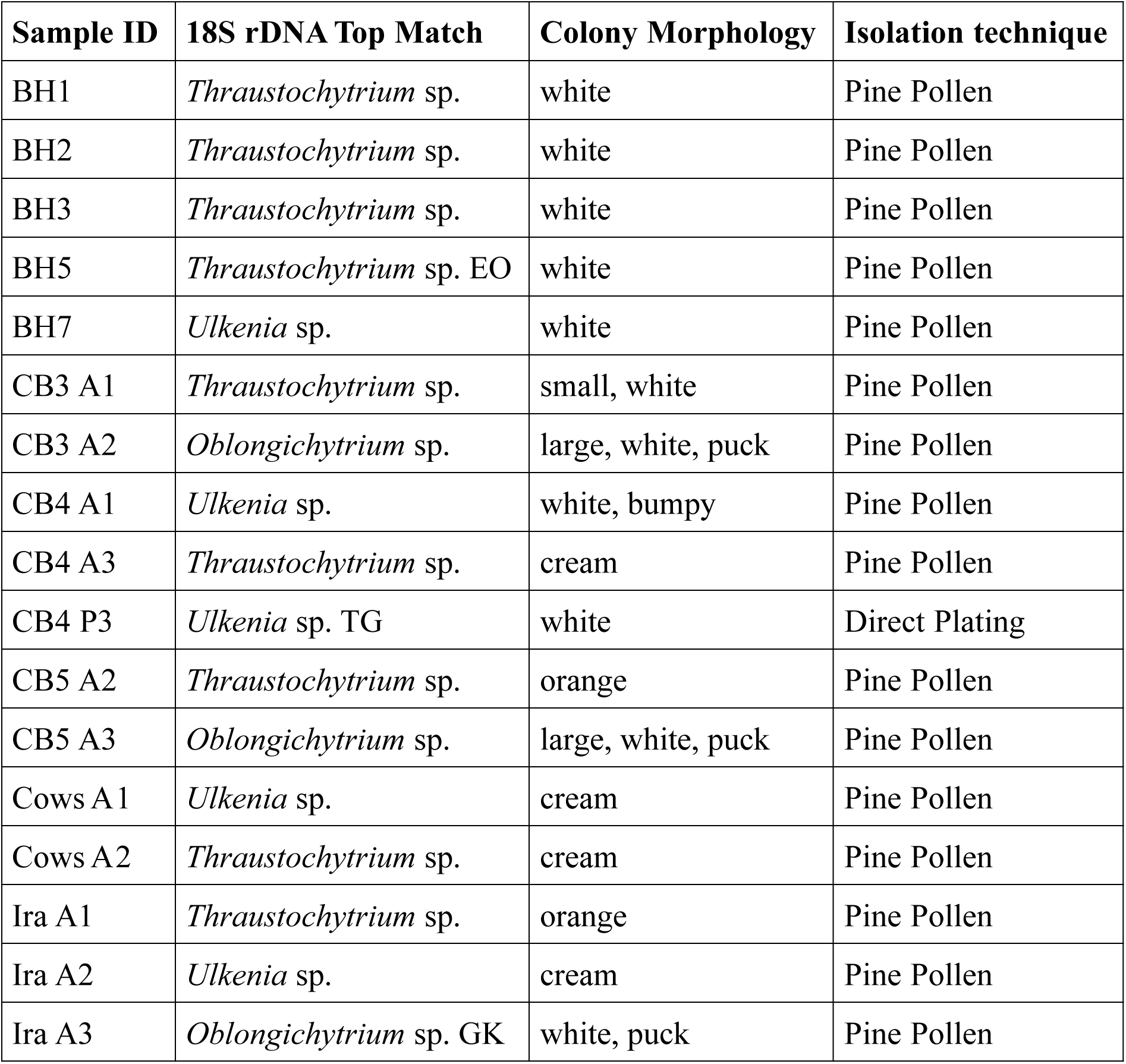
Description of 17 cultured Labyrinthulomycetes from Nova Scotian sampling sites. Each sample ID corresponds to the site of origin (Table 1) with the genus determined from the best 18S rDNA BLAST hit, brief description of colony morphology on 790 By+ plates and the isolation technique (pine pollen baiting or direct plating).

| **Sample ID** | **18S rDNA Top Match** | **Colony Morphology** | **Isolation technique** |
| --- | --- | --- | --- |
| BH1 | *Thraustochytrium* sp. | white | Pine Pollen |
| BH2 | *Thraustochytrium* sp. | white | Pine Pollen |
| BH3 | *Thraustochytrium* sp. | white | Pine Pollen |
| BH5 | *Thraustochytrium* sp. EO | white | Pine Pollen |
| BH7 | *Ulkenia* sp. | white | Pine Pollen |
| CB3 A1 | *Thraustochytrium* sp. | small, white | Pine Pollen |
| CB3 A2 | *Oblongichytrium* sp. | large, white, puck | Pine Pollen |
| CB4 A1 | *Ulkenia* sp. | white, bumpy | Pine Pollen |
| CB4 A3 | *Thraustochytrium* sp. | cream | Pine Pollen |
| CB4 P3 | *Ulkenia* sp. TG | white | Direct Plating |
| CB5 A2 | *Thraustochytrium* sp. | orange | Pine Pollen |
| CB5 A3 | *Oblongichytrium* sp. | large, white, puck | Pine Pollen |
| Cows A1 | *Ulkenia* sp. | cream | Pine Pollen |
| Cows A2 | *Thraustochytrium* sp. | cream | Pine Pollen |
| Ira A1 | *Thraustochytrium* sp. | orange | Pine Pollen |
| Ira A2 | *Ulkenia* sp. | cream | Pine Pollen |
| Ira A3 | *Oblongichytrium* sp. GK | white, puck | Pine Pollen |

The partial 18S rDNA sequences determined to belong to labyrinthulomycetes were subjected to phylogenetic analysis. Our maximum likelihood phylogenetic tree (**Figure 2**) places members of the genus *Oblongichytrium* as the basal branch of the Labyrinthulomycetes, consistent with previous studies (Tsui et al. 2009; Pan et al. 2017). However, we did not include environmental clades (e.g., LAB12-16) (Pan et al. 2017) or sequences from *Amphitremida* or *Amphifilidae*, as neither morphology or preliminary BLAST searches suggested any isolates belonged to these clades (Pan et al. 2017; Takahashi et al. 2014; Tice et al. 2016; Anderson and Cavalier-Smith 2012). Notably, none of our new isolates belong to the genera *Aurantiochytrium* or *Schizochytrium.* Prior studies have produced many strains of *Aurantiochytrium* from plant detritus (Wang et al. 2019; Damare 2014) likely due to ease of culturing and their wide distribution. In Nova Scotia, we found no *Aurantiochytrium* or *Schizochytrium* strains in the three sites examined, despite previous reports in the Atlantic region (Burja et al. 2006).

**Figure 2.**
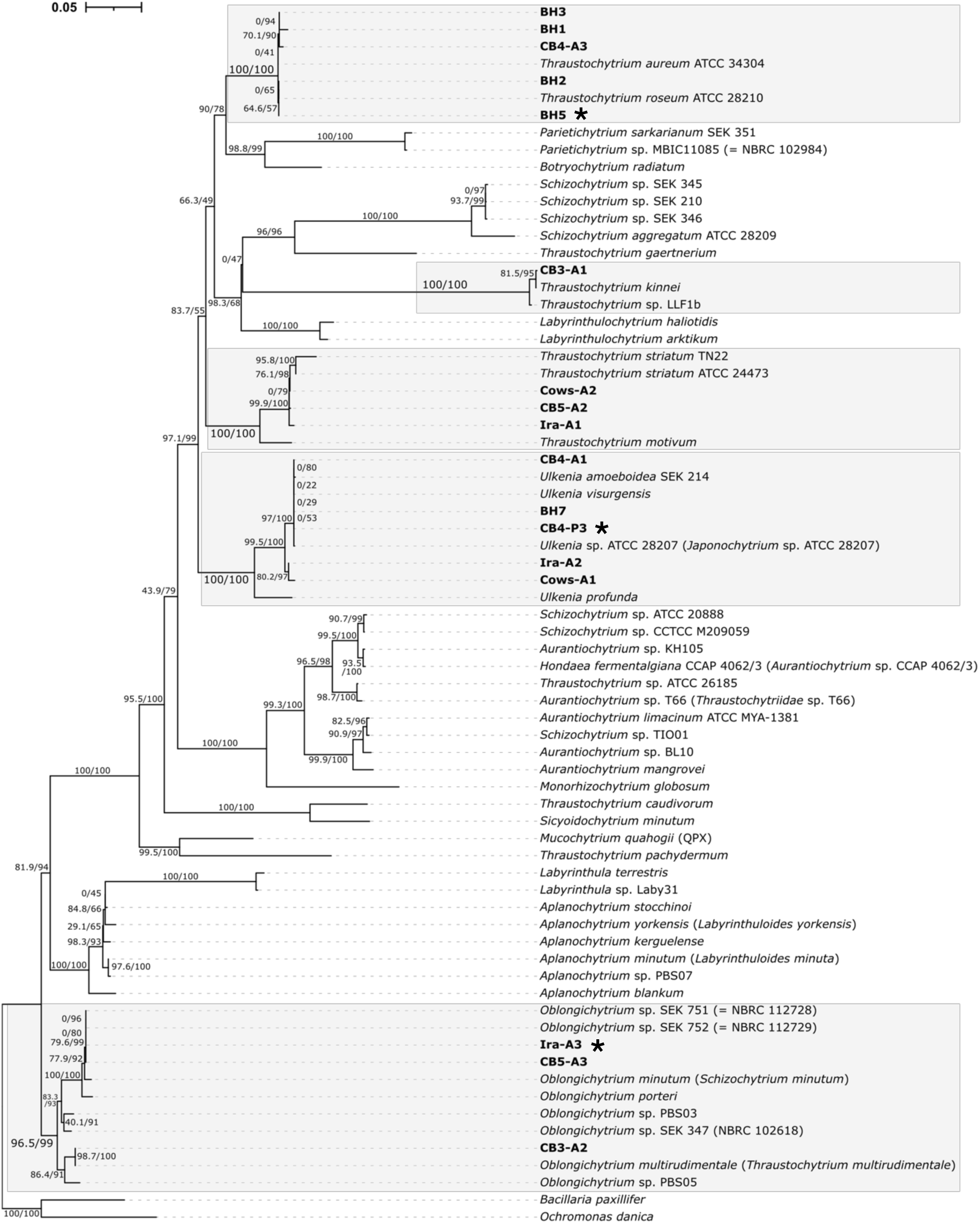
18S rDNA phylogenetic tree of Labyrinthulomycetes from SILVA SSU and environmental isolate. Shown is a consensus IQ-Tree maximum-likelihood tree (model TN+F+I+R3 selected by IQ-Tree using ModelFinder plus; branch support is percentage of 1000 aLRT/UFboot replicates). Sequences from newly obtained environmental isolates are shown in bold and stars indicate strains characterized in this study. Scale bar corresponds to inferred number of nucleotide substitutions per site.

Phylogenetic analysis suggests that three of our isolates (Ira-A3, CB5-A3 and CB3-A2) are species of *Oblongichytrium*. Ira-A3 and CB5-A3 cluster with *Oblongichytrium minutum* with high support (aLRT/UFboot = 100/100) suggesting they are part of clade Ob1 (Ueda et al. 2015) while CB3-A2 is highly similar to *Oblongichytrium multirudimentale* (99.88 % identity across 1710 nucleotides) suggesting it is part of clade Ob2 (Ueda et al. 2015). Unfortunately, our CB3-A2 culture perished after a few passages, which has been reported previously for labyrinthulomycetes isolates (Collado-Mercado et al. 2010).

Three of our 18S rDNA sequences (CB4-A1, BH7 and CB4-P3) were found to cluster very closely with *Ulkenia* species with high support (aLRT/UFboot = 97/100); these three sequences are highly similar and cannot be distinguished from each other or to *Ulkenia* spp. sequences based on 18S rDNA phylogeny alone. The Ira-A2 and Cows-A1 isolates form their own clade (aLRT/UFboot = 80.2/97) within the genus *Ulkenia*, distinct from type strains such as *U. virgensis*, *U. amoeboidea* and *U. profunda*. There are no available genomes for members of the genus *Ulkenia*, making these new environmental isolates compelling candidates to understand labyrinthulomycetes genome diversity.

The remaining nine strains are all members of *Thraustochytrium* (Figure 2). It should be noted that this genus has been determined to be a paraphyletic entity, with a handful of recognized subclades named after type species: *T. aureum*, *T. striatum* and *T. kinnei* (Pan et al. 2017; Ueda et al. 2015). Five of our strains (BH1, BH2, BH3, BH5 and CB4-A3) branch robustly with the *T. aureum* group (aLRT/UFboot = 100/100), while three are *T. striatum* (Ira-A3, Cows-A2 and CB5-A3; aLRT/UFboot = 99.9/100). Lastly, strain CB3-A1 is highly similar to *T. kinnei* (98.4% identity over 1,723 nucleotides; aLRT/UFboot = 100/100).

18S rDNA phylogenetic analysis revealed that labyrinthulomycetes strains isolated from three distinct locations in Nova Scotia belong to very different genera, exhibiting with- and between-site taxonomic diversity. With the goal of further characterizing our new isolates, one strain from each of the genera we identified was further studied by transmission electron microscopy (TEM), differential interference contrast (DIC) and fluorescence microscopy to characterize their ultrastructure.

### Light and transmission electron microscopy

#### Thraustochytrium sp. EO

Isolate BH5 was identified as a strain of *Thraustochytrium aureum* and tentatively named *Thraustochytrium* sp. EO. This strain was isolated from an estuary region northeast of Windsor, Nova Scotia and shares high 18S rDNA identity (99.6% over 826 nucleotides) with the type strain *T. roseum* ATCC 28210, a close relative to *T. aureum* ATCC 34304. Like *T. aureum*, *Thraustochytrium* sp. EO has globose cells and, under the growth conditions used herein, minimally developed ectoplasmic nets. In liquid culture these cells form colonies made of tens of vegetative cells and sporangia (**Figure 3A, 3B**). Both DNA staining and TEM show individual vegetative cells of *Thraustochytrium* sp. EO to be multinucleate (**Figure 3C-E**). Prominent ultrastructural features of this strain include the thraustochytrid-defining bothrosome, scales, lipid bodies, electron dense regions and Golgi apparatus (**Figure 3F, 3G**).

**Figure 3.**
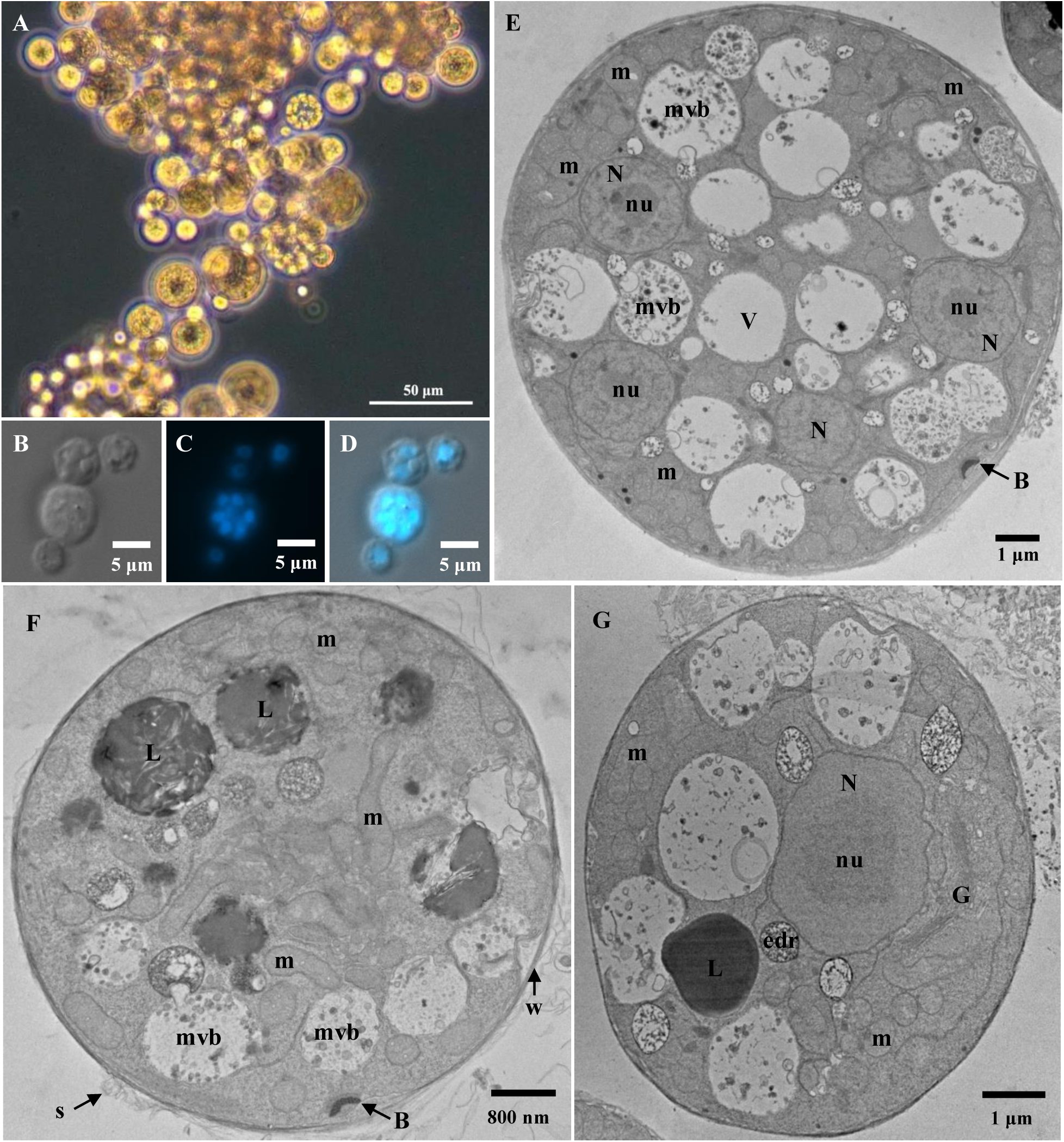
Micrographs of *Thraustochytrium* sp. EO. **A.** Brightfield light micrograph of cell clusters in liquid 790 By+ cultures. **B.** DIC micrograph **C.** Hoechst 33342 staining. **D.** Merged image of B and C. **E-G.** Transmission electron micrographs. Labelled structures are nucleus (N), nucleolus (nu), mitochondria (m), bothrosome (B), scales (S), lipid droplet (L), cell wall (w), electron dense regions (edr), Golgi apparatus (G), and multivesicular body (mvb).

#### Ulkenia sp. TG

CB4-P3, the only labyrinthulomycete successfully isolated by direct plating, was established from a sample collected at St. Ann’s provincial park in Cape Breton, Nova Scotia. This isolate has its 18S rDNA best BLAST hit to *Japonochytrium* sp. ATCC 28207 (e-value 0.0, identity 97.6%), a strain that is now recognized as a species of *Ulkenia* (Yokoyama et al. 2007). Given this nomenclatural amendment and the high support (aLRT/UFboot = 97/100) for both CB4-P3 and *Japonochytrium* sp. ATCC 28207 as members of the *Ulkenia* clade (**Figure 2**), we have designated the CB4-P3 isolate *Ulkenia* sp. TG. *Ulkenia* sp. TG has globose and subglobose cells and like previous descriptions of the genus, disperse and form smaller aggregations compared to other genera (Yokoyama et al. 2007) (**Figure 4A**). Large sporangia can be seen with TEM (**Figure 4B**) indicating *Ulkenia* sp. TG forms large sporangia similar to those described as aplanospores in *U. visurgensis* (Moss 1980). Similarly, we found cells of this strain to have variable diameter and numbers of nuclei depending on the life stage (**Figure 4 C-E**). TEM revealed mono- (**Figure 4 F**) or multi-nucleate (**Figure 4 G**) cells as well as characteristic ultrastructure such as Golgi, electron dense regions, lipid droplets and a bothrosome (**Figure 4 F-H**).

**Figure 4.**
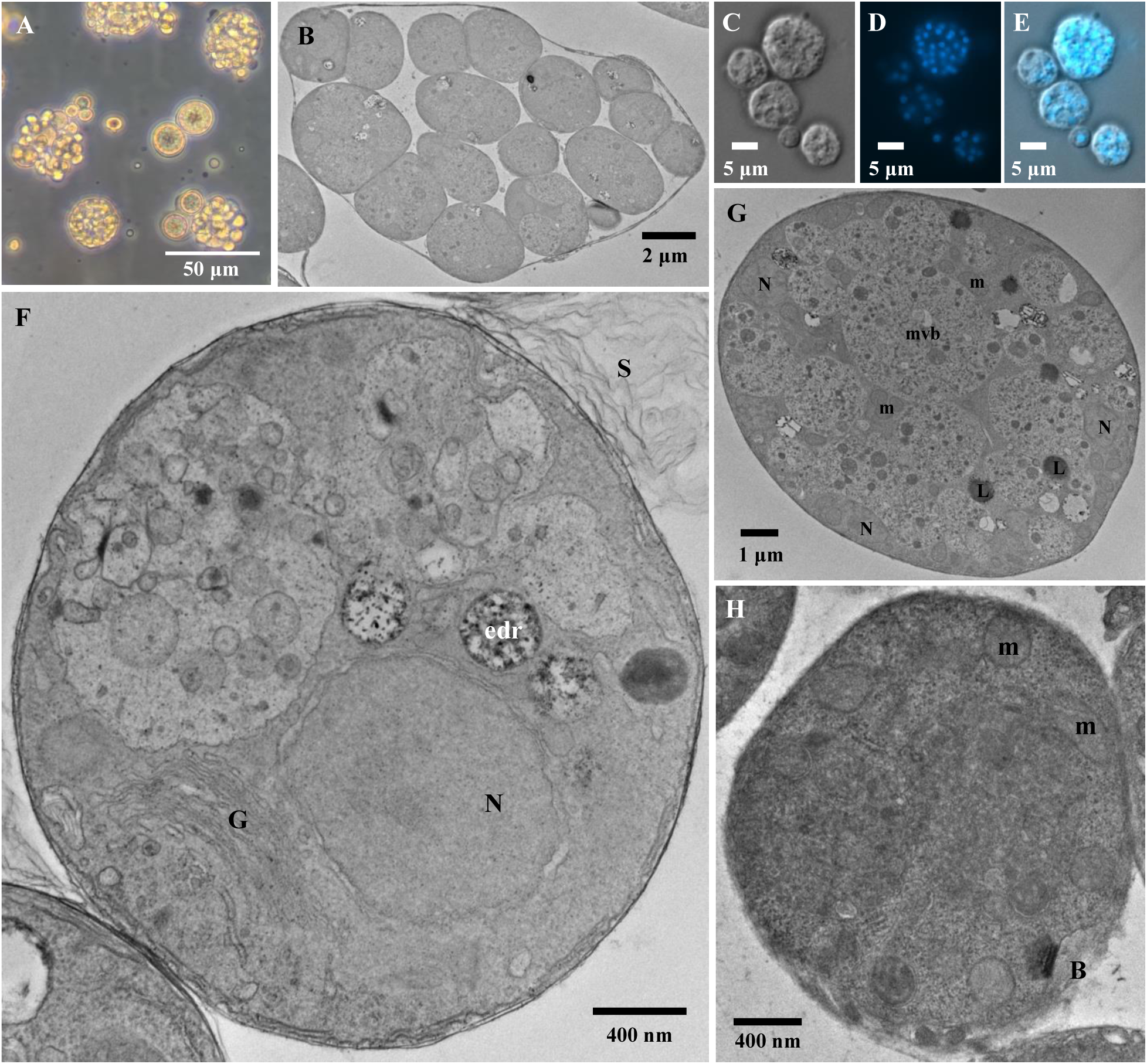
Micrographs of *Ulkenia* sp. TG **A.** Light micrograph of cell clusters in liquid culture. **B.** TEM of sporangium **C.** DIC micrograph **D.** Hoechst 33342 staining. **E.** Merged image of panel C and D. **F-H.** Transmission electron micrographs. Labelled structures are nucleus (N),, mitochondria (m), bothrosome (B), lipid droplet (L), electron dense regions (edr), scales (S), Golgi apparatus (G), and multivesicular body (mvb).

#### Oblongichytrium sp. GK

The isolate Ira-A3 shares high 18S rDNA identity with *Oblongichytrium* sp. SEK 752 (98.1%) and *Oblongichytrium* sp. SEK 708 (98.1%). As noted, this strain clusters strongly (aLRT/UFboot = 100/100) with *Oblongichytrium* spp. SEK 752 and SEK 708, as well as *O. minutum* (basionym: *Schizochytrium minutum*) (**Figure 2**). The genus *Oblongichytrium* was established by Yokoyama et al. in 2007 as part of the taxonomic rearrangement of *Schizochytrium* (Yokoyama and Honda 2007). Many of the morphological characteristics are similar between *Oblongichytrium* and *Schizochytrium*, but 18S rDNA phylogeny resolves distinct clades, as shown here (**Figure 2**) and elsewhere (Pan et al. 2017; Tice et al. 2016; Collado-Mercado et al. 2010; Yokoyama and Honda 2007). *Oblongichytrium* sp. GK has well-developed ectoplasmic nets under fed conditions (**Figure 5A, 5B**) and forms large colonies containing tens to hundreds of cells in liquid culture (**Figure 5A**). The vegetative cells are frequently seen to be multinucleate in DIC-DAPI overlay as the colonies divide by continuous binary cell division within the cell wall (**Figure 5C-E**). TEM also identified recently divided cells with two plasma membrane-delineated cells within the cell wall (**Figure 5F**). Mature cells are approximately 5 μm across with well-defined nuclei, Golgi and mitochondria (**Figure 5G**), while the oblong or elliptical shaped zoospores are smaller with flagella reminiscent of strains like *Oblongichytrium* sp. SEK 347 (**Figure 5H**) (Yokoyama and Honda 2007). While the type species *O. minutum* (basionym: *Schizochytrium minutum*) (Gaertner 1981) and *O. multirudimentale* (basionym: *Thraustochytrium multirudimentale*) (Goldstein 1963) have been described with light microscopy, we are only aware of *Oblongichytrium* sp. SEK 347 TEM micrographs (Yokoyama and Honda 2007). The TEM data presented here for an *Oblongichytrium* aff. *minutum* strain thus add to key ultrastructural information for this poorly studied genus.

**Figure 5.**
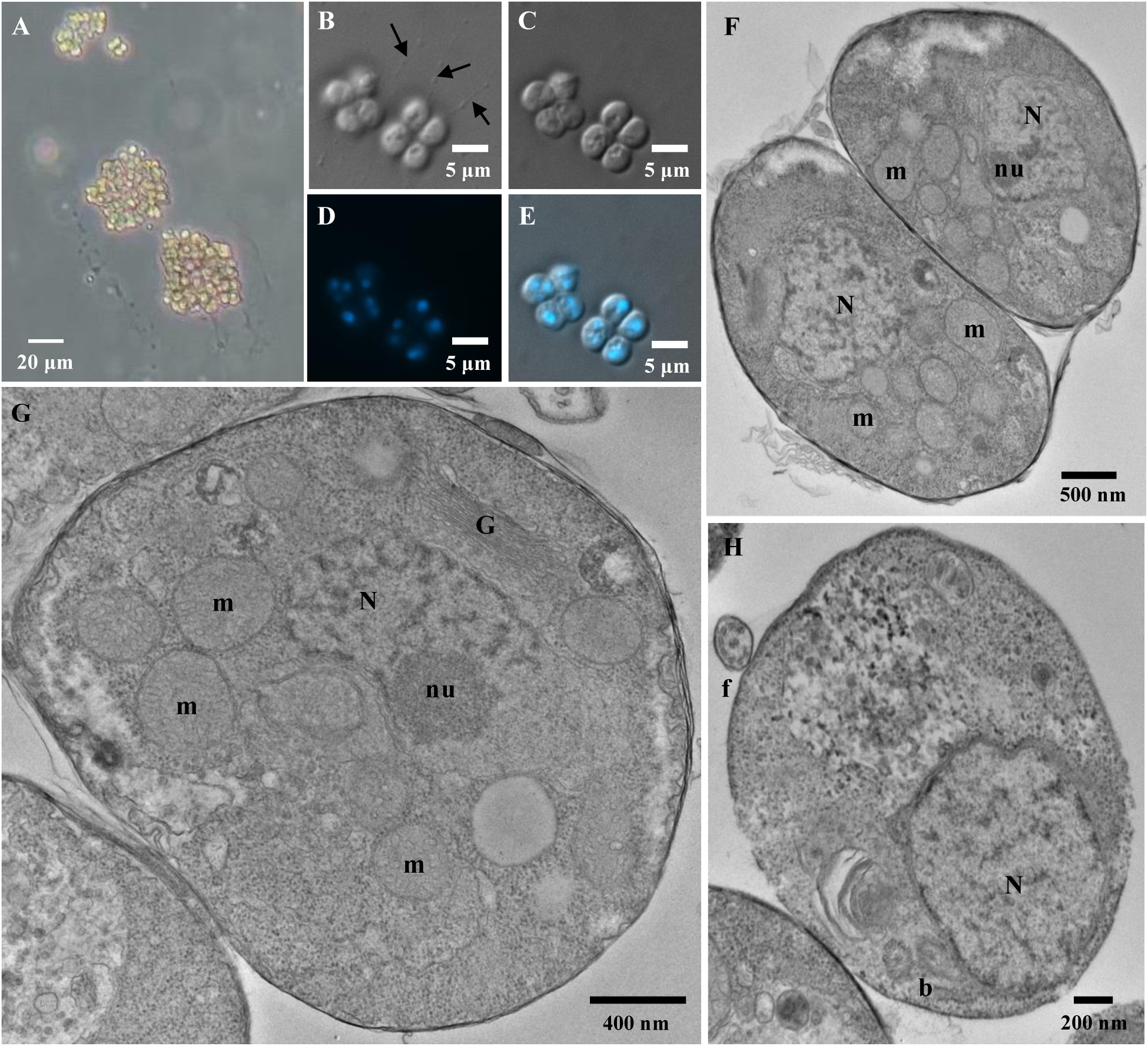
Micrographs of *Oblongichytrium* sp. GK **A.** Brightfield light micrograph of cell clusters in liquid 790 By+ cultures. **B.** DIC light micrograph showing ectoplasmic nets (arrows). **C.** DIC micrograph of panel B with cell bodies in focus. **C.** DAPI staining of panel C. **D.** Merged image of panel C and D. **F-H.** Transmission electron micrographs. Labelled structures are nucleus (N), nucleolus (nu), mitochondria (m),, Golgi apparatus (G), flagellum (f), and basal body (b) .

## CONCLUSION

Much of the previous research on Labyrinthulomycetes has focused on environmental sequence diversity and cultivation or growth optimization of high omega-3 producing strains for biotechnological applications. Consequently, clades such as *Oblongichytrium* and *Ulkenia* have been neglected and type strains like *O. minutum* have been lost or are no longer offered from culture collections. We have isolated several novel environmental strains of labyrinthulomycetes from Atlantic Canada, expanding the known genetic diversity of thraustochytrids and enabling further molecular investigation of these ubiquitous marine protists. Of note is the potential for these new strains to form the foundation of a broad comparative genomic investigation into the biology and ecology of these organisms. The evolution of thraustochytrids has recently become of interest because *Aurantiochytrium liminacium* has been shown to be host to mirusviruses (Chung et al. 2025; Collier et al. 2023), a new lineage of large DNA viruses found in globally in marine environments (Gaïa et al. 2023), The genetic signatures of mirusviruses have been detected in diverse lineages across the eukaryotic tree (Zhao et al. 2024); within Bigyra, the distribution of mirusviruses within Labyrinthulomycetes is unclear. Genome sequencing of the strains established herein should make it possible to address this question; having the cultures themselves will facilitate dissection of host-virus interactions.

## ACKNOWLEDGMENTS

We thank Mary Ann Trevors at the Electron Microscopy Core Facility (Dalhousie University) for preparing samples for transmission electron microscopy, as well as training and technical support. We also thank Gerard Gaspard at the Cellular and Molecular Digital Imaging Facility (Dalhousie University) for training and technical support for operating the DIC and florescence microscope. Additional thanks to Alastair Simpson for facilitating sampling at the BH site. JL was the recipient of NSERC CGS-M and CGS-D (601146). This research was funded by a Discovery Grant from the Natural Sciences and Engineering Research Council of Canada (RGPIN-2019-05058) awarded to JMA.

